# Rats show a preference for certain unfamiliar strains of rats

**DOI:** 10.1101/2021.02.18.431764

**Authors:** Hiroki Kogo, Yasushi Kiyokawa, Yukari Takeuchi

## Abstract

Humans show distinct social behaviours when we recognise social similarity in opponents that are members of the same social group. However, little attention has been paid to the role of social similarity in non-human animals. In Wistar subject rats, the presence of an unfamiliar Wistar rat mitigated stress responses, suggesting the importance of social similarity in this phenomenon. We found that the presence of unfamiliar Sprague-Dawley (SD) or Long-Evans (LE) rats, but not an unfamiliar Fischer 344 (F344) rat, similarly mitigated stress in subject rats. It is therefore possible that the subject rats recognised social similarity to unfamiliar SD and LE rats. In this study, we demonstrated that Wistar subject rats were capable of categorizing unfamiliar rats based on their strain, and that Wistar subjects showed a preference for unfamiliar Wistar, SD, and LE rats over F344 rats. However, the subject rats did not show a preference among Wistar, SD, and LE rats. In addition, the results were not due to an aversion to F344 rats, and preference was not affected when anaesthetised rats were presented to subject rats. The findings suggested that rats recognise social similarity to certain unfamiliar strains of rats.

## Background

In social animals, behaviours towards conspecific individuals are influenced by two independent factors. One is individual familiarity with an opponent, which is assessed based on the individual identity of the opponent. The clearest examples of such behaviours would include those observed exclusively between a mother and her infant or between pair-bonded mates [1, 2]. Behaviours towards conspecific individuals other than mothers or mates are also affected by individual familiarity; for example, animals are prone to exhibit pro-social behaviour towards familiar individuals [3], and antagonistic behaviour (e.g., defending territory) towards unfamiliar individuals [4]. The other factor that affects behaviours towards conspecific individuals is social similarity to the opponent. In such cases, when an animal evaluates an opponent as being a member of the same group (i.e., ingroup), then the animal recognises social similarity to the opponent, regardless of individual familiarity with the opponent. The significance of social similarity and its effect on behaviour have been studied extensively in humans. For example, even people who explicitly endorse egalitarian values are typically not free of biases related to the race or ethnicity of an opponent [5, 6]. Nonetheless, relatively few studies have investigated the role that social similarity plays in interactions among non-human animals.

Social buffering refers to the phenomenon in which the presence of affiliative conspecific individuals mitigates stress responses in a subject [7]. Studies have demonstrated that, in addition to social buffering by a mother or mate [8, 9], social buffering can be elicited by other conspecific individuals in a variety of species [10–12]. We conducted a series of studies in rats to examine whether social buffering is induced by a conspecific individual other than the subject’s mother or mate [13, 14]. Stress responses caused by an aversively conditioned stimulus were completely blocked when a Wistar subject rat was accompanied by an unfamiliar same-sex Wistar rat, hereafter referred to as the accompanying rat [15–19]. Therefore, social similarity alone appears to be sufficient for inducing social buffering in rats.

In addition to a number of characteristics [20–24], including the importance of olfactory signals [20, 25, 26] and possible neural mechanisms [26–29], social buffering in rats was found to be induced by certain unfamiliar strains of unfamiliar individuals. Specifically, an unfamiliar Sprague-Dawley (SD), Long-Evans (LE), or Lewis accompanying rat had the effect of acting as a social buffer and induced social buffering in the Wistar subject [30]. Conversely, unfamiliar Fischer 344 (F344) or Brown Norway accompanying rats were not regarded as buffers, hereafter referred to as non-buffers, because they did not induce social buffering [30]. Given that the dyad had not met before, individual familiarity with the accompanying rat could not contribute towards social buffering. The remaining possibility might be that the Wistar subject recognised some social similarity to the accompanying rat, even if the subject was unfamiliar with the SD, LE, and Lewis strains. Examining the genealogy of these strains reveals that Lewis rats are derived from Wistar rats, and SD and LE rats are three quarters and half Wistar rats, respectively. Conversely, F344 and Brown Norway rats were established independently of Wistar rats [31]. Therefore, SD, LE and Lewis rats are relatively closely related to Wistar rats, although specific information on actual genetic distances is not currently available. It has been demonstrated that humans will show a preference for an unfamiliar individual if they consider that there is some level of social similarity with that individual [6]. Based on these findings, we hypothesised that a subject would show preference for social buffers over non-buffers.

In the present study, a series of experiments was performed using Wistar rats as subjects. Before testing the hypothesis, in Experiment 1, we first examined whether the rats were capable of categorizing unfamiliar rats based on their strain. In a habituation-dishabituation test, the subject was sequentially exposed to three unfamiliar Wistar rats and one unfamiliar F344 rat. In Experiment 2, we examined whether the subject showed a preference for social buffers over non-buffers. We prepared unfamiliar Wistar, SD, and LE rats as social buffers, and unfamiliar F344 rats as non-buffers. In the partner preference test, the subject was offered a choice between a social buffer and non-buffer. The choices between the subject’s own strain of social buffer and other unfamiliar strains of social buffer were also offered in order to assess if the subject showed a preference among social buffers. In Experiment 3, we assessed if the subject showed an aversion to non-buffers. The subject was offered a choice between being alone or spending time with a social buffer or non-buffer. In Experiment 4, we assessed whether differences in the behaviour of social buffers and non-buffers contributed to the preference of the subject. The subject was also offered the choice of an anaesthetised social buffer or an anaesthetised non-buffer.

## Methods

All experiments were approved by the Animal Care and Use Committee of the Faculty of Agriculture at The University of Tokyo, according to guidelines adapted from the *Consensus Recommendations on Effective Institutional Animal Care and Use Committees* by the Scientists Center for Animal Welfare.

### Animals

Experimentally naïve male Wistar (aged 7 weeks), SD (aged 7 weeks), LE (aged 7 weeks), and F344 (aged 9 weeks) rats were purchased from Charles River Laboratories Japan (Kanagawa, Japan). To ensure that body size was similar among rats, we ordered F344 rats of different ages [29, 30]. Upon arrival, all rats were housed individually in a room with an ambient temperature of 24 ± 1°C, humidity of 45 ± 5%, and a 12-h light/12-h dark cycle (lights were switched on at 8:00). Food and water were administered *ad libitum*. Nylon zip ties (Panduit Japan, Tokyo, Japan) were used as colour bands and attached to the stimulus rats more than 4 h after arrival, mostly on the following day. All behavioural tests were performed during the light period of the day after the attachment of the colour band.

### Procedures

#### Experiment 1

The habituation-dishabituation test was performed in an illuminated room using a two-chamber apparatus. The apparatus consisted of two parallel polycarbonate chambers (standard rat cage: 24 × 40 × 20 cm). A stainless tube (6 cm diameter, 8 cm long) connected the two chambers at approximately one-third the length of the long side of the chambers. One chamber was defined as the starting chamber while the other was defined as the stimulus chamber. The subject was first placed in the starting chamber and allowed to explore the entire apparatus for 10 min. Then, a stimulus rat was tethered in the stimulus chamber to restrict its movement within the chamber. The behaviour of the subject during the subsequent 10-min test period was recorded with a video camera (DCR-TRV18; Sony, Tokyo, Japan) and an HDD-BD recorder (DMR-BW770; Panasonic, Osaka, Japan). After the test period, the subject was returned to the colony room and kept undisturbed for 10 min. The same procedure was repeated until the subject (*n* = 6) was sequentially presented with three unfamiliar Wistar stimulus rats (individuals A, B, and C) and one unfamiliar F344 stimulus rat (individual D).

#### Experiment 2

The partner preference test was performed in an illuminated room using a three-chamber apparatus that was prepared by connecting one identical chamber to the two-chamber apparatus used in Experiment 1. Specifically, three parallel polycarbonate chambers were linearly connected by stainless tubes located at one-third the length of the long side of the chambers. The centre chamber was defined as the starting chamber. The chambers at both ends of the apparatus were defined as the stimulus chambers. The subject was first placed in the starting chamber and allowed to explore the entire apparatus for 20 min. Then, an unfamiliar Wistar, SD, LE, or F344 stimulus rat was tethered individually in each stimulus chamber to restrict its movement within the cage. Each stimulus rat was tethered on opposite sides of the apparatus on consecutive runs. The 60-min test period was started when the subject had visited both stimulus chambers more than once and then returned to the starting chamber. During the test period, the subject was allowed to explore a pair of Wistar and F344 stimulus chambers (*n* = 7), a pair of SD and F344 stimulus chambers (*n* = 8), a pair of LE and F344 stimulus chambers (*n* = 7), a pair of Wistar and SD stimulus chambers (*n* = 8), or a pair of Wistar and LE stimulus chambers (*n* = 8). The behaviour of the subject was recorded with a video camera (DCR-TRV18; Sony, Tokyo, Japan) and an HDD-BD recorder (DMR-BW770; Panasonic, Osaka, Japan).

#### Experiment 3

The partner preference test was performed as described in Experiment 2 with the exception that only one unfamiliar stimulus rat was used for each subject. Accordingly, the chamber in each end of the apparatus was defined as the empty and stimulus chamber, respectively. During the test period, the subject was allowed to explore a pair of empty and Wistar stimulus chambers (*n* = 7), a pair of empty and SD stimulus chambers (*n* = 6), a pair of empty and LE stimulus chambers (*n* = 7), or a pair of empty and F344 stimulus chambers (*n* = 6).

#### Experiment 4

The partner preference test was performed as described in Experiment 2 with the exception that the stimulus rats were anaesthetised with pentobarbital sodium (54 mg/kg, i.p.). During the test period, the subject was allowed to explore a pair of anaesthetised Wistar and anaesthetised F344 stimulus chambers (*n* = 8).

### Data analyses and statistical procedures

A researcher recorded the time spent in each chamber (the centre of the forepaws was considered to indicate the location of the subject) and the time engaged in active social interaction with the stimulus rat (the subject touched the stimulus rat with its nose and/or allo-groomed the stimulus rat). Microsoft Excel-based Visual Basic software was used to record the duration of key presses, as in our previous studies [32, 33]. The data were analysed using the paired *t* test. The significance level was set at *P* < 0.05 for all statistical tests.

## Results

Data are expressed as mean ± standard error of the means (SEM).

### Experiment 1

We first assessed if the subject could categorise unfamiliar rats based on their strain. In this experiment, the subject was sequentially presented three unfamiliar Wistar stimulus rats (individuals A, B, and C) and one unfamiliar F344 stimulus rat (individual D) in the habituation-dishabituation test (Fig. 1). The time engaged in social interactions with the third Wistar stimulus rat (individual C) was shorter than that with the first Wistar stimulus rat (individual A) (*t_5_* = −5.14, *P* < 0.01). In addition, the time engaged in interactions with the first F344 stimulus rat (individual D) was longer than that with the third Wistar stimulus rat (individual C) (*t*_5_ = 5.47, *P* < 0.01). In contrast, the time spent in the stimulus chamber did not differ between when the first or third Wistar stimulus rat was presented (*t*_5_ = −1.89, *P* = 0.12) (Table 1). Similarly, the time spent in the stimulus chamber did not change when the third Wistar stimulus rat was replaced with the first F344 stimulus rat (*t*_5_ = 1.35, *P* = 0.24) (Table 1). These results suggested that the subject could categorise unfamiliar rats based on their strain.

**Fig. 1.**
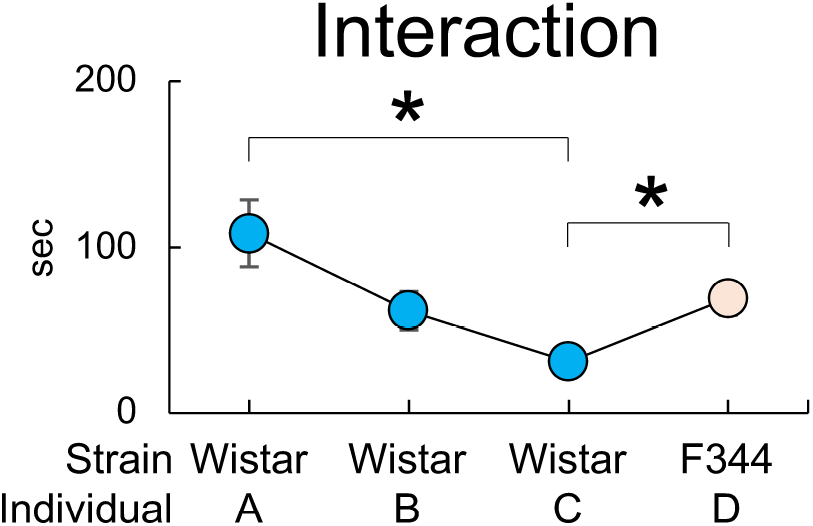
Time the subject (*n* = 6) engaged in social interaction with three unfamiliar, sequentially presented, Wistar stimulus rats (individuals A, B, and C) and one unfamiliar Fischer 344 (F344) stimulus rat (individual D) in Experiment 1. \**P* < 0.05 with paired *t*-test.

**Table 1.** Time spent in the stimulus chamber in Experiment 1

| Stimulus rat | Time spent in the chamber (s) |
| --- | --- |
| Wistar (individual A) | 509 ± 17 |
| Wistar (individual B) | 429 ± 18 |
| Wistar (individual C) | 448 ± 30 |
| F344 (individual D) | 470 ± 30 |

Data are expressed as mean ± standard error of the mean.
F344: Fischer 344.

### Experiment 2

We assessed whether the subject showed a preference for social buffers over non-buffers in the partner preference test (Fig. 2). When a pair of Wistar and F344 stimulus rats, a pair of SD and F344 stimulus rats, or a pair of LE and F344 stimulus rats was presented to the subject, the time engaged in social interactions was longer with the Wistar (*t*_6_ = −3.71, *P* < 0.05), SD (*t*_7_ = −2.43, *P* < 0.05), or LE stimulus rats (*t*_6_ = −3.20, *P* < 0.05) than with the F344 stimulus rat. When we analysed the time spent in each chamber, the time spent in the Wistar (*t*_6_ = −3.80, *P* < 0.01), SD (*t*_7_ = −2.47, *P* < 0.05), and LE stimulus chambers (*t*_6_ = −3.58, *P* < 0.05) was longer than that spent in the F344 stimulus chamber (Table 2). These results suggested that the subject showed a preference for social buffers over non-buffers.

**Fig. 2.**
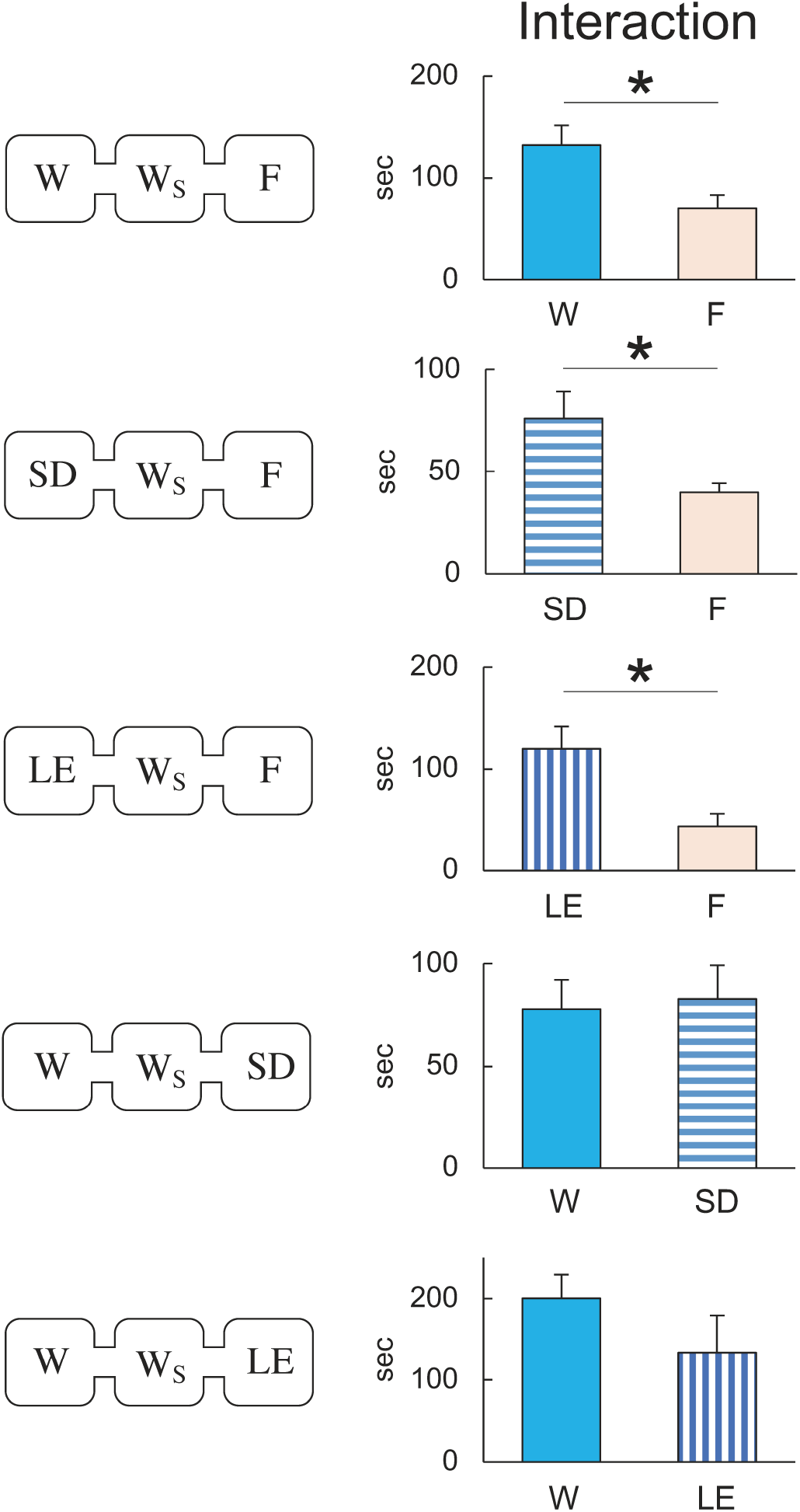
Time the Wistar subject (W_S_) engaged in social interaction with the stimulus rats when allowed to explore a pair of Wistar (W) and Fischer 344 (F) stimulus chambers (*n* = 7), a pair of Sprague-Dawley (SD) and F stimulus chambers (*n* = 8), a pair of Long-Evans (LE) and F stimulus chambers (*n* = 7), a pair of W and SD stimulus chambers (*n* = 8), or a pair of W and LE stimulus chambers (*n* = 8) in Experiment 2. \**P* < 0.05 with paired *t*-test.

**Table 2.** Time spent in the stimulus chamber in Experiment 2

| Stimulus rat | Time spent in the chamber (min) |
| --- | --- |
| Wistar vs. F344 |  |
| Wistar | 28.3 ± 2.3* |
| F344 | 13.8 ± 2.1 |
| SD vs. F344 |  |
| SD | 27.5 ± 4.0* |
| F344 | 14.0 ± 2.1 |
| LE vs. F344 |  |
| LE | 23.3 ± 3.2* |
| F344 | 8.8 ± 1.7 |
| Wistar vs. SD |  |
| Wistar | 24.4 ± 3.6 |
| SD | 17.0 ± 2.5 |
| Wistar vs. LE |  |
| Wistar | 28.4 ± 5.3 |
| LE | 13.8 ± 4.7 |

Data are expressed as mean ± standard error of the mean.
F344: Fischer 344; LE: Long-Evans; SD: Sprague-Dawley.
\* $P < 0.05$ for paired $t$ -test.

We also assessed whether the subject showed a preference between social buffers. When a pair of Wistar and SD stimulus rats or a pair of Wistar and LE stimulus rats was presented to the subject rat, the time engaged in social interactions with the Wistar stimulus rat was similar to that with the SD (*t*_7_ = 0.35, *P* = 0.74) or LE stimulus rat (*t*_7_ = −1.14, *P* = 0.29), respectively (Fig. 2). Similarly, the time spent in the Wistar stimulus chamber was the same as that in the SD (*t*_7_ = −1.66, *P* = 0.14) or LE stimulus chamber (*t*_7_ = −1.52, *P* = 0.17) (Table 2). These results suggested that the subject rats did not show a preference among between buffers.

### Experiment 3

Since the partner preference test compares the relative preference intensity of the two stimulus rats, it is unclear whether the subject showed a preference for social buffers or showed aversion to non-buffers. To clarify this point, the subject was offered a choice between being alone and spending time with a social buffer or a non-buffer. Since there was no stimulus rat in the empty cage, we could not measure the time engaged in social interactions with the stimulus rat in the empty cage. However, we confirmed that the subject interacted with all strains of stimulus rat (Table 4). When we analysed the time spent in each chamber, the time spent in the Wistar (*t*_6_ = −5.57, *P* < 0.01), SD (*t*_5_ = −5.64, *P* < 0.01), LE (*t*_6_ = −3.73, *P* < 0.01), or F344 stimulus chamber (*t*_5_ = −6.73, *P* < 0.01) was longer than it was in the empty chamber (Table 3). These results suggested that the subject did not show an aversion to non-buffers.

**Table 3.** Results of Experiment 3

| Stimulus rat | Time spent in the chamber<br>(min) | Time engaged in interaction<br>(s) |
| --- | --- | --- |
| Wistar vs. Empty |  |  |
| Wistar | 32.4 ± 3.6* | 453 ± 78 |
| Empty | 6.8 ± 1.4 | N/A |
| SD vs. Empty |  |  |
| SD | 34.6 ± 4.5* | 141 ± 23 |
| Empty | 6.1 ± 0.9 | N/A |
| LE vs. Empty |  |  |
| LE | 33.0 ± 5.0* | 213 ± 34 |
| Empty | 7.2 ± 2.5 | N/A |
| F344 vs. Empty |  |  |
| F344 | 37.0 ± 4.3* | 93.3 ± 12.5 |
| Empty | 4.2 ± 0.7 | N/A |

Data are expressed as mean ± standard error of the mean.
F344: Fischer 344; LE: Long-Evans; N/A: not applicable; SD: Sprague-Dawley.
\* $P < 0.05$ for paired $t$ -test.

**Table 4.**
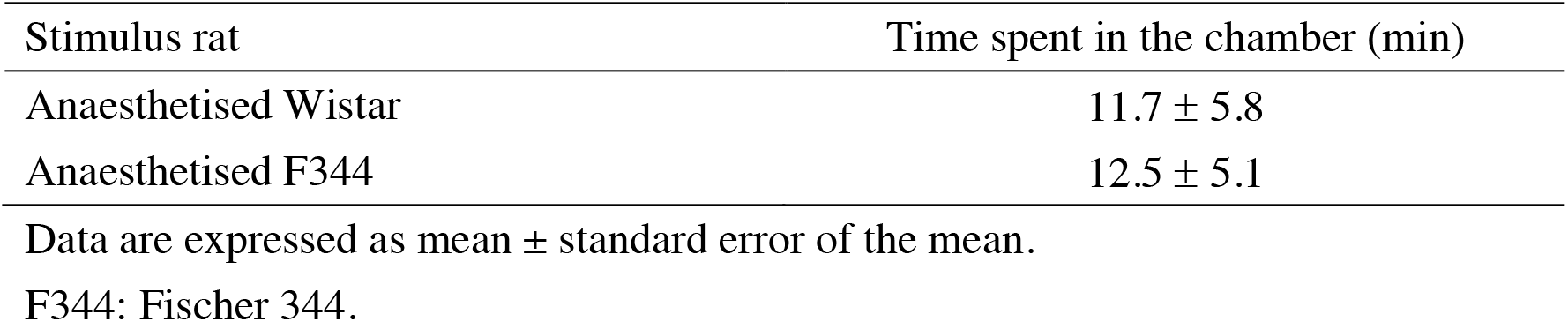
Time spent in the stimulus chamber in Experiment 4

### Experiment 4

It is known that F344 rats are less responsive to social interactions with Wistar rats. For example, co-housing a Wistar rat with an F344 rat is an experimental model of peer-rejection in Wistar rats [34–36]. It is therefore possible that the behaviour of stimulus rats may determine the preference of the subject. To clarify this point, the subject was offered a choice between an anaesthetised social buffer of the same strain as the subject and an anaesthetised non-buffer (Fig. 3). The time engaged in active interaction was longer with the anaesthetised Wistar stimulus rat than with the anaesthetised F344 stimulus rat (*t*_7_ = −2.48, *P* < 0.05). However, the time spent in the chamber was similar between the two chambers (*t*_7_ = 0.10, *P* = 0.92) (Table 4). These results suggested that the subject’s preference was less affected by the behaviour of the stimulus rat.

**Fig. 3.**
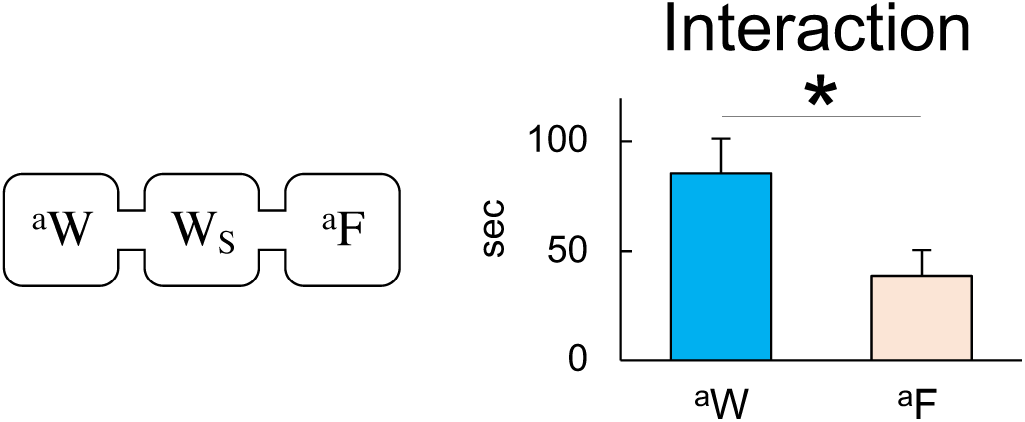
Time the Wistar subject (W_S_) engaged in social interaction with the stimulus rats when allowed to explore a pair of anaesthetised Wistar (^a^W) and anaesthetised Fischer 344 (^a^F) stimulus chambers (*n* = 8) in Experiment 4. \**P* < 0.05 with paired *t*-test.

## Discussion

In Experiment 1, the subject showed habituation after the sequential presentation of three unfamiliar Wistar rats. The subject then showed dishabituation when an unfamiliar F344 rat was presented as the fourth individual. These results suggested that the subject could categorise unfamiliar rats based on their strain. In Experiment 2, the subject showed a preference for social buffers over non-buffers. In addition, the subject did not show a preference between social buffers. These results could not be ascribed to the alternative interpretation that the subject showed aversion to non-buffers, because the subject showed a preference for non-buffers over being alone in Experiment 3. In addition, the results of Experiment 2 were not likely due to differences in behaviour between social buffers and non-buffers, because the subject showed a preference for anaesthetised social buffers over anaesthetised non-buffers in Experiment 4. Based on these results, we concluded that the subject recognised social similarity to certain unfamiliar strains of rats.

The results of Experiment 1 provided the first evidence that rats can categorise unfamiliar rats based on their strain. Given the importance of olfactory signals in the recognition of other rats [37], it is possible that the subject recognised a signature mixture of each strain. A signature mixture is a subset of the molecules in an animal’s chemical profile that is used to recognise an animal as a member of a particular group [38, 39]. Body odours, one of the sources of the signature mixture, are determined by genetic factors [40–42]. Because individuals of inbred strains are genetically identical, most components of their body odour are thought to be shared among individuals and to serve as a signature of the inbred strain. Even among individuals of outbred strains, several genetic factors are known to be shared. For example, Wistar rats shared the same allele at 6 out of 27 genetic loci examined among individuals [43]. Therefore, it is reasonable to assume that some components of their body odour are also shared among individuals and that they contribute to the signature mixture of the outbred strain. Taken together, it is possible that the unique odour mixture of each strain enabled the subject to categorise unfamiliar rats based on their strain.

A series of partner preference tests demonstrated that the subject showed a preference for social buffers over non-buffers. Based on these findings, we propose that rats do not consider all unfamiliar strains to be equivalent. Several studies have employed unfamiliar rat strains as opponents in order to assess the role of social similarity in social learning and/or expression of prosocial behaviour [44–46]. However, all of the studies selected the unfamiliar strain based simply on the fact that the subject rat was unfamiliar with the opponent strain. The findings of the present study suggest that the Wistar subject rats classified unfamiliar SD and LE rats as ingroup members. Consistent with this classification is the observation in a previous study in which pairings of unfamiliar SD and LE rats both induced vicarious fear responses in the SD subject [45]. Therefore, it is more appropriate for researchers to select unfamiliar strains of rats based on genealogy or genetic distances, if available. However, in a previous study, the SD subject did not release an unfamiliar LE rat from a restrainer [44]. If the classification suggested in the present study was indeed employed by subject rats, then the SD subject rats should have classified the unfamiliar LE rats as ingroup members and showed helping behaviour. One possible reason for this discrepancy might be the differences in the position of the subject. In the first two studies, the subject was in a position where it could benefit from the opponent, such as receiving buffering effects or danger information. In contrast, the subject had to make an effort to help the restrained rat in the last study. In such a situation, it is reasonable to assume that the subject evaluated the opponent strictly and classified the opponent rats with a narrower range of genetic background as ingroup members. Further analyses are necessary to understand how rats classify opponents as being ingroup members.

## Conclusion

In the present study, we demonstrated that the subject showed a preference for social buffers over non-buffers. These results suggest that rats recognise social similarity to certain unfamiliar strains of rats. The preference observed in the present study shares characteristics with ingroup favouritism in humans. Ingroup favouritism describes the phenomenon in which a person shows a preference for a member of the same group. Race or ethnicity is known to be one of the salient factors that automatically induces such ingroup favouritism [47]. In addition, ingroup favouritism and outgroup derogation are not necessarily reciprocally related [6, 48]. Given that ingroup favouritism based on race and ethnicity emerges in the early stages of life [49], it seems to be a biologically important phenomenon in social animals. Therefore, further analyses on preference among unfamiliar strains of rats may shed light on the neurobiology of sociality in animals.

